# Nav1.7 gating in human iPSC derived sensory neurons: an experimental and computational study

**DOI:** 10.1101/2020.08.04.235861

**Authors:** Alberto Capurro, Jack Thornton, Bruno Cessac, Lyle Armstrong, Evelyne Sernagor

## Abstract

Chronic pain is a global healthcare problem with a huge societal impact. Its management remains unsatisfactory, with no single treatment clinically approved in most cases. In this study we use an *in vitro* experimental model of erythromelalgia consisting of sensory neurons derived from human induced pluripotent stem cells obtained from a patient (carrying the mutation F1449V) and a control subject. We combine neurophysiology and computational modelling to focus on the Nav1.7 voltage gated sodium channel, which acts as an amplifier of the receptor potential in nociceptive neurons and plays a critical role in erythromelalgia due to gain of function mutations causing the channel to open with smaller depolarisations.

Using multi-electrode array (extracellular) recordings, we found that the scorpion toxin OD1 increases the excitability of sensory neurons in cultures obtained from the control donor, evidenced by increased spontaneous spike rate and amplitude. In erythromelalgia cultures, the application of the Nav1.7 blocker PF-05089771 effectively stopped spontaneous firing. These results, which are in accordance with current clamp and voltage clamp recordings reported in the literature, are explained with a conductance-based computational model of a single human nociceptive neuron. The disease was simulated through a decrease of the Nav1.7 half activation voltage, which decreased the rheobase and increased the response to supra threshold depolarizing currents. This enhanced response could be successfully supressed by blocking the Nav1.7 channels. The painful effects of OD1 were simulated through a slower establishment and a quicker removal of Nav1.7 inactivation, reproducing the effects of the toxin on the spike frequency and amplitude. Our model simulations suggest that the increase in extracellular spike amplitude observed in the MEA after OD1 treatment can be due mainly to a slope increase in the ascending phase of the intracellular spike caused by impaired inactivation gating.

## Introduction

Chronic pain is a global healthcare problem, particularly affecting elderly people, women and persons with lower socio-economic status (Van Hecke et al., 2013). It is one of the most common reasons for physician consultation in developed countries, interfering with quality of life and causing large socio-economic impacts that include significant loss of working hours and the need of clinical care. Current therapies have limitations in their effectiveness and side effects, creating an urgent need to develop more precise and effective treatments for pain management (Khouzam, 2000). In this study we focus on the voltage dependent properties of the sodium channel Nav1.7 that is implicated in inherited erythromelalgia, a rare chronic condition causing attacks of severe pain (e.g., McDonnell et al., 2016).

Pain evoked spiking activity starts in peripheral terminals of dorsal root ganglion (DRG) neurons. The central extensions of these neurons form the Aδ and C fibres which establish glutamatergic synapses onto second order neurons within the spinal cord. They can be stimulated by mechanical, thermal or chemical stimuli as well as by inflammatory mediators. Distributed throughout the body (skin, viscera, muscles, joints, meninges), they carry noxious sensory information into the central nervous system (Serpell, 2006). The recent discovery of nociceptive Schwann cells has changed the notion of bare nerve terminals being the starting point of pain sensation, and introduced the concept of a glio-neural end organ in the skin that transmits nociceptive information to the nerve, resembling the specialized receptor cells found in other sensory systems (Abdo et al., 2019).

DRG neurons express several types of voltage gated sodium channels with different properties (Rush et al., 2007; Lera Ruiz and Kraus, 2015). The opening of Nav1.7 channels requires smaller depolarisations than other types of sodium channels, being more easily activated by the graded generator potentials. For this reason it is considered to be a threshold channel (Dib-Hajj et al., 2013) that acts as an amplifier of the receptor potential, increasing the probability of triggering a spike via the activation of other higher threshold sodium channels such as Nav1.8 (e.g., Renganathan, 2001; Payne, et. al., 2015). This amplification property makes Nav1.7 a major contributor to pain signalling in humans.

Erythromelalgia is caused by gain‑of‑function mutations of the gene SCN9A, which encodes for the Nav1.7 channels (Dib-Hajj et al., 2005; McDonnell et al., 2016). Different types of mutations have in common the need for less depolarization to open the channel than in the wild type, resulting in a decrease of the rheobase (i.e., minimal amplitude of a depolarizing current of infinite duration that is able to evoke a spike). Neurons often fire spontaneously in erythromelelgia cultures, while control cells are silent unless receiving strong depolarizing currents (Cao et al., 2016).

In this study, we have used extracellular recordings of spontaneous activity in sensory neuronal cultures derived from human induced pluripotent stem cells (hiPSCs) obtained from an erythromelalgia patient and a control subject to study the voltage dependent gating properties of Nav1.7 channels. To start, the effects on the neuronal firing of two different pharmacological compounds-a pain eliciting scorpion toxin (OD1, Motin et al., 2016) and a potentially analgesic drug (PF-05089771, Cao et al., 2016)-were assessed in these cultures using multi electrode array (MEA) recordings. We then presented a simple conductance-based computational model of a single nociceptive neuron to explain our findings in terms of the Nav1.7 gating process.

## Materials and Methods

### Cells and MEA recordings

For this study we used hiPSCs from a control subject (cell line AD3) and an erythromelalgia patient (cell line RCi002-A, carrying the mutation F1449V) made available at the European Bank for induced pluripotent stem cells (EBiSC). The cells were differentiated into sensory neurons using a small molecule based protocol described previously (Cao et al., 2016, and references therein). Once they reached this stage, the cells were re-plated in 24 wells MEA plates (MEA700, Multichannel Systems, Reutlingen, Germany) where the spontaneous activity was recorded after 10 weeks of maturation. Each well contains 12 circular electrodes (100 µm diameter, 700 µm electrode pitch), making a total of 288 electrodes per plate. The large distance between electrodes makes it very unlikely that the activity originating from an individual neuron will be detected by adjacent electrodes. On the other hand, due to the relatively large cell density in these cultures, which organize in structures called neural rosettes (e.g., Wilson and Stice, 2006), it is quite likely that one electrode can detect the activity of more than one neuron (up to four in our data sets, although most channels had only one or two).

We compared the spontaneous activity of sensory neurons obtained from a control subject (plated in 18 wells, making a total of 216 electrodes in MEA 1) with an erythromelalgia patient (plated in 24 wells, making a total of 288 electrodes in MEA 2). Each well constitutes an independent cell culture.

The cultures were treated with OD1 (100 nM) in the control subject (MEA 1) and with PF-05089771 (100 nM) in the patient (MEA 2). The activity was recorded for 5 minutes immediately before the application of each substance and compared with a recording of the same duration performed 5 minutes after the onset of drug exposure. Wells contained 200 µl of growing medium and 5 µl drops were added to apply the treatments. The experimental doses were selected to be in a near saturation range to ensure a strong effect, based on dose response curves published previously (Cao et al., 2016 for PF-05089771; Motin et al., 2016 for OD1). Both compounds were purchased from Tocris (Bio-Techne, Abingdon, UK).

### Spike sorting and pairing of units

Recordings were performed with the Multiwell-MEA-System and the software Multi Channel Experimenter (Multi Channel Systems, Reutlingen, Germany). Each channel was band-pass filtered (100 to 3500 Hz) and acquired with a sampling rate of 20 kHz.

The raw voltage traces were first plotted and inspected using a zoom tool to discard artefacts and confirm the existence of spikes. The voltage time series were then fed into the Matlab toolbox Waveclus (Quian Quiroga et al., 2004) to perform spike sorting in all active channels. In the toolbox, the continuous data were filtered again with a non-causal band pass filter between 300 and 3000 Hz and the firing times were detected with an amplitude threshold. We used a dual threshold (i.e., picking deflections in both up and down directions) set to 5 median absolute deviations of the filtered voltage signal with a refractory period of 2 ms to avoid double annotations due to fast voltage oscillations in the vicinity of the threshold. Spikes were aligned to their maximum, after interpolating the waveforms to locate the peak time more accurately. The toolbox uses a wavelet based method for feature extraction, and the grouping of spikes into clusters is done with super-paramagnetic clustering (Blatt et al., 1996), a stochastic algorithm that does not assume any particular distribution of the data. In few cases the dual threshold created mistakes in the detection, so we decided to use a single threshold, kept for the units that corresponded to the same neuron before and after a given treatment. We consider this spike sorting strategy as supervised, always following the criterion of the biologist as ground true. The firing times and voltage cut outs of all units were stored to document the parameters and quality of the spike sorting in each channel.

Units that were active both before and after each pharmacological treatment were identified. The only set-in-stone criterion for deciding if they are the same cell is the coordinates of the current sources, using high density MEAs (e.g., Hilgen et al., 2017). As our data were recorded with low density MEAs, we cannot provide this level of certainty, but assumed that they corresponded to the same neuron if the wave shape remained similar across recordings performed within few minutes of each other. Possible ambiguities were minimized by the fact that most channels yielded only one or two active neurons. These pairs were used to investigate the changes in spike frequency and amplitude caused by the pharmacological treatments (Figs 1 and 2). If we were reasonably convinced that a given unit corresponded to the same neuron before and after treatment, we designated the two recordings as a pair. In the case of the control cultures, few neurons had spontaneous discharges and most cells started to fire only after the OD1 treatment. Conversely, in the case of the patient cultures, several neurons were active before, but few remained active after the application of PF-05089771.

**Fig 1.**
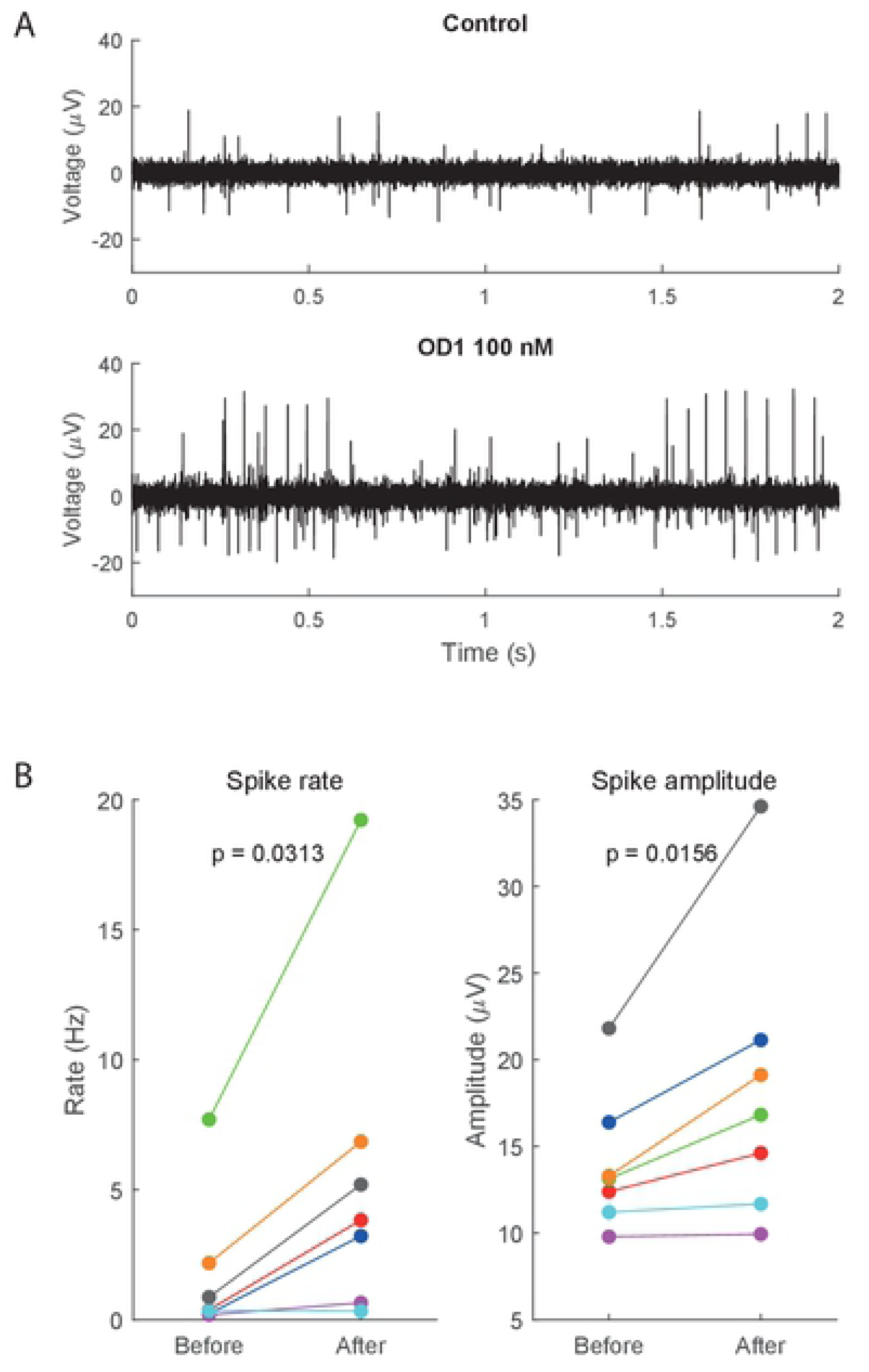
Effects of OD1 in hiPSC derived sensory neurons. (A) Representative example of voltage data in MEA 1 before (upper panel) and after (lower panel) OD1 (100 nM) application. Note the increase in rate and amplitude of the spontaneous spikes. (B) Rate and amplitude of paired neurons before and after OD1 (100 nM) application. Corresponding pairs in both plots are colour coded. Wilcoxon signed rank test p values are printed in each panel. The normalized changes of both indexes are highly correlated (r = 0.78).

**Fig 2.**
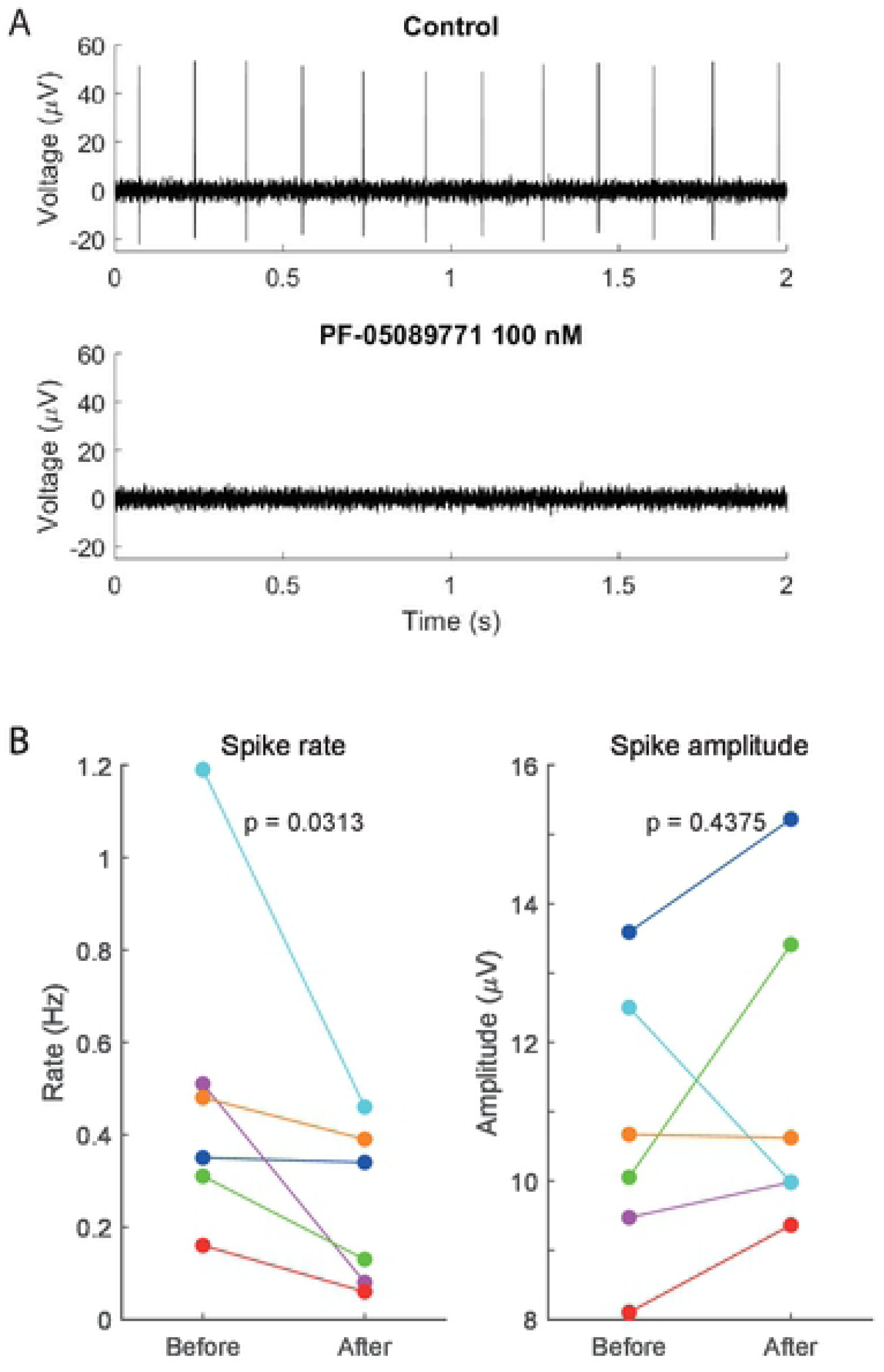
Effects of PF05089771 in hiPSC derived sensory neurons. (A) Representative example of voltage data before and after PF-05089771 (100 nM). Note the firing suppression. (B) Rate and amplitude of the neurons paired before and after treatment. The normalized changes in both indexes are poorly correlated (r = 0.21).

For statistical comparisons between two conditions we used the Wilcoxon signed rank test (for paired values) or the Mann–Whitney U test (for unpaired values). We also used cross correlation histograms of the spike times recorded from the same channel to assess whether neurons tend to fire at fixed delay from another.

### Numerical simulations

We performed numerical simulations of the membrane potential and sodium channels gate parameters of a single nociceptive neuron using the software NEURON (www.neuron.yale.edu) and the Channel Builder GUI (McDougal et al., 2017). Following previous models of DRG neurons (Kovalsky et al., 2009; Choi and Waxman, 2011; Vasylyev et al., 2014; Verma et al., 2020), we used Hodgkin and Huxley (HH) type of differential equations, which have been shown to be relevant in the context of the mammalian nervous system (Krouchev et al., 2017). To keep the model as simple as possible, only a single current (Nav1.7) was added to the original HH scheme. The two different types of sodium channels included differed only in their parameters (Table 1), the form of the equations was the same (Eq. 1 to 8 in S1 Appendix). Integration method was the IDA algorithm (e.g., Carnevale, 2007) with a fixed time step (*dt*) of 0.0025 ms (i.e., 400 time steps per ms). This very small *dt* value was selected to obtain smooth plots when focusing on the brief time window (e.g., 15 ms) surrounding a single spike (Fig. 4).

**Table 1.**
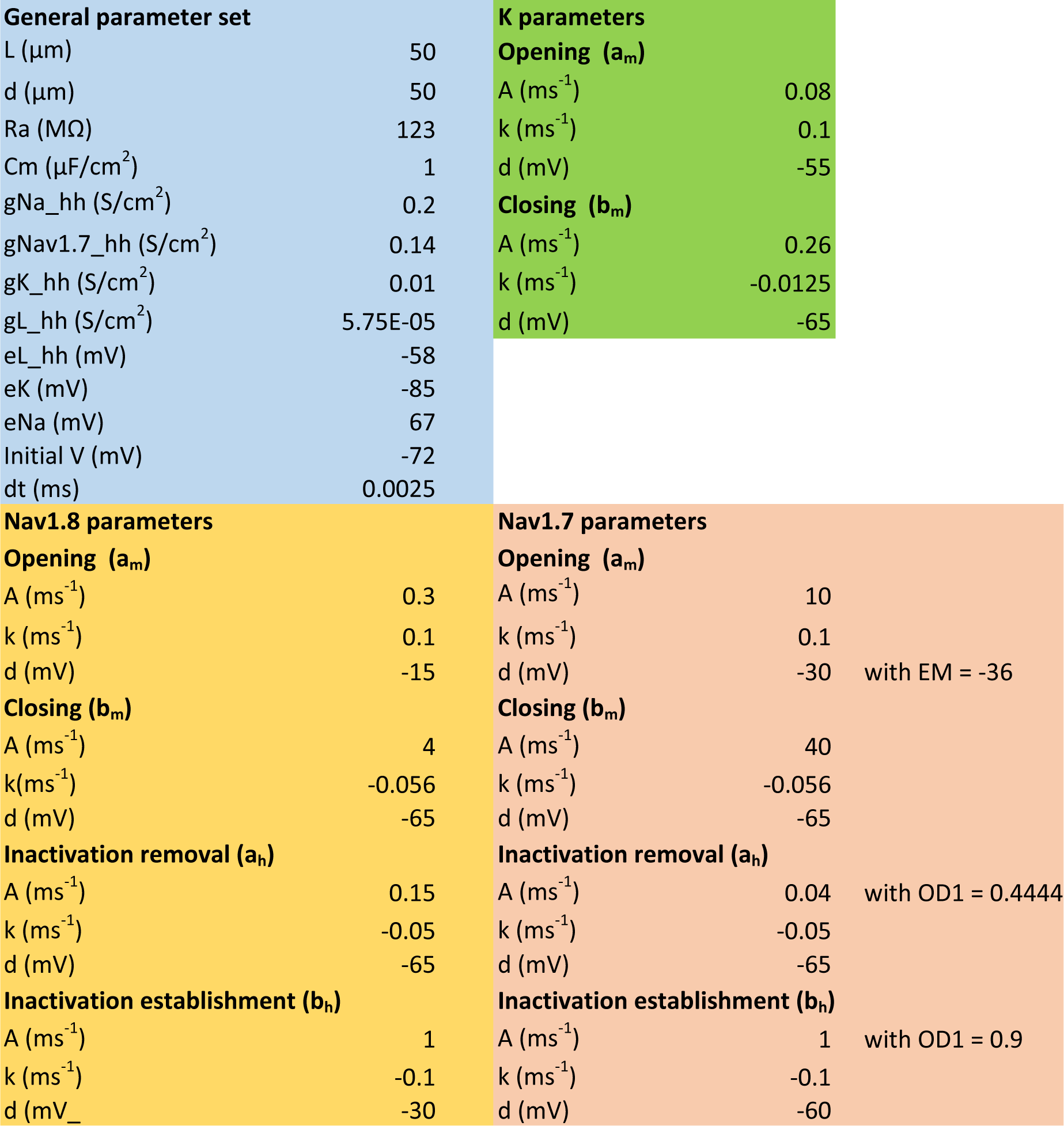
Default parameter values used for the simulations. Particular cases are indicated in the corresponding figures.

The parameters (Table 1) are plausible values obtained from rat and human experiments that appeared in the literature. The equilibrium potentials of the ions, as well as the leak conductance and cytosolic resistance, were from the modelling study of Choi and Waxman (2011), based on whole cell patch-clamp recordings performed in rat DRG neurons (Choi et al., 2007). For the size/capacitance of the cell we relied on human studies (Zhang et al., 2017), which reported that DRG neurons are moderately bigger than in rat and that the maximum conductance of the ions is generally higher in human cells. We set our soma as a 50 µm diameter and height cylinder, corresponding to a small human DRG cell in the rat/human comparison histogram presented in the figure 2 of Zhang et al. (2017). The proportion of *g_maxNa_* values between both Na^+^ currents was as reported in Choi and Waxman (2011), with Nav1.7 reaching roughly 0.7 of Nav1.8 value, although we used larger *g_max_* values for all currents. Using this *g_maxNa_* proportion, the electric charge underlying the spike is mainly provided by Nav1.8, as it has been reported in C-type DRG neurons (Renganathan et al., 2001).

The half activation and half inactivation voltages of the sodium currents were graphically estimated from the study of Rush et al. (2007) performed in rat DRG neurons (see figure 1–D and E in Rush et al., 2007), assuming that the TTX sensitive component of the current corresponds to Nav1.7. The study of Motin et al. (2016) shows a similar voltage-dependence profile of Nav1.7 activation and inactivation (see figure 3–B in Motin et al., 2016, done with human Nav1.7 channels introduced in CHO cells), despite the difference in experimental preparations. The main differences between our representation of both Na^+^ currents (Table 1) are the following. Nav1.8 had larger conductance (0.2 S/cm^2^) and more a depolarized half activation (*d_m_* = −15 mV) and half inactivation voltage (*d_h_* = −30 mV). Nav1.7 had half activation and half inactivation voltages shifted in the hyperpolarizing direction (*d_m_*= −30 mV, *d_h_* = −60 mV) and a smaller value of maximum conductance (0.14 S/cm^2^). We also followed literature reports stating that Nav1.7 has faster opening/closing rates and slower recovery from inactivation (repriming) than Nav1.8 (Dib-Hajj et al., 2013; Dib Hamaed, 2019). These gating properties were implemented through the corresponding A values (Table 1), that multiply the voltage dependent rates (Eq. 4, 5, 7 and 8). Slow repriming is a relevant feature because it enables the low frequency firing typical of C fibres (Dib-Hajj et al., 2013). Close state inactivation (Herzog et al., 2003) was not implemented in the present model, as it would require a more complex Markov-type kinetic (e.g., Andreozzi et al., 2019).

**Fig 3.**
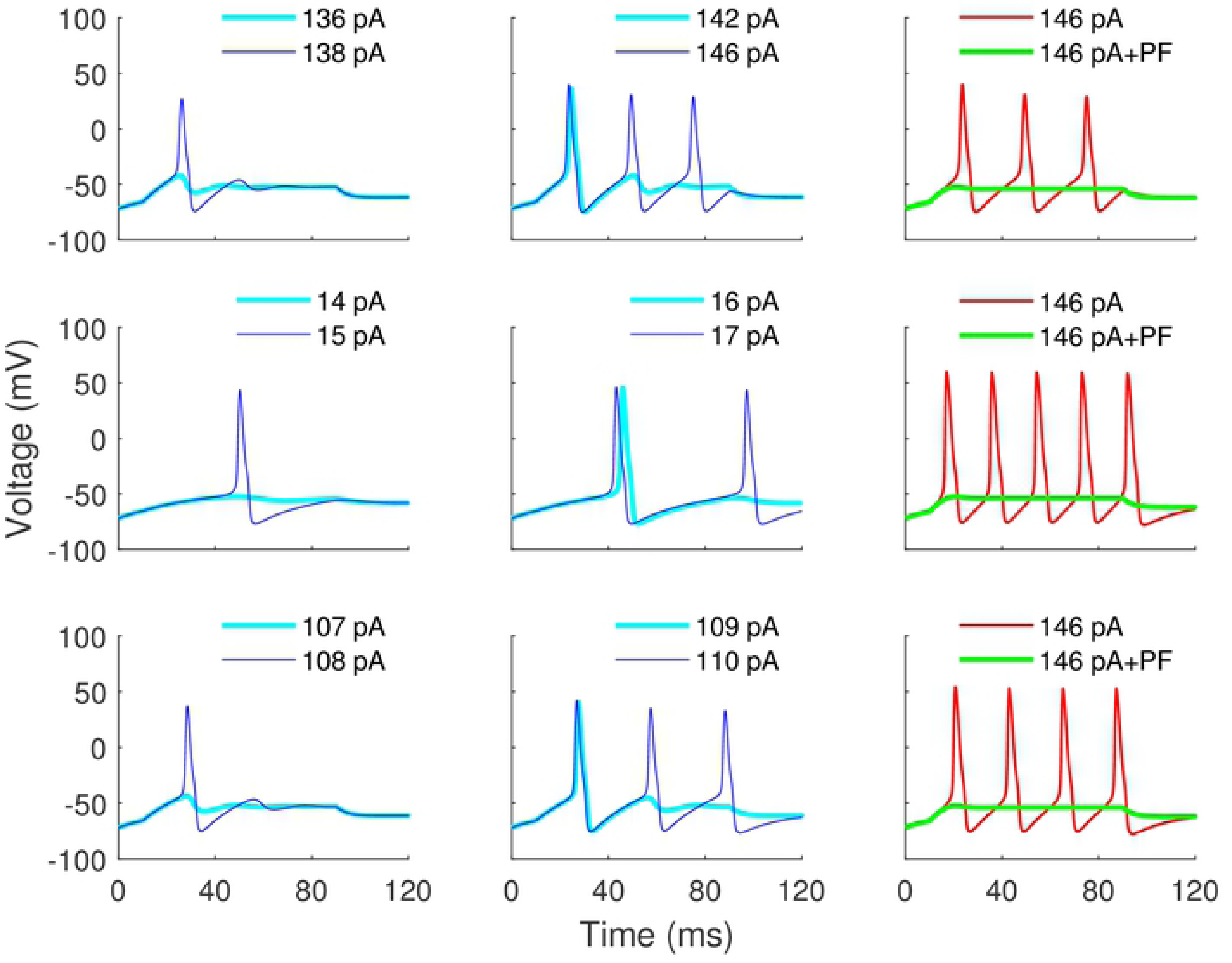
Simulations of membrane potential. Control conditions (upper row), erythromelalgia (middle row) and OD1 application (bottom row). The left column shows rheobases and the middle column shows thresholds for sustained spiking. Supra-threshold responses and PF-05089771 treatment are depicted in the right column.

To simulate erythromelalgia we varied the half activation voltage of Nav1.7 (*d* in Eq. 4) from −30 to −36 mV. This shift is within the range reported for the F1449V mutation (Dib-Hajj et al., 2005). PF-05089771 effect at a saturating dose was simulated by setting the maximum conductance of Nav1.7 (g maxNa in Eg. 2) to 0. In the case of OD1, we simultaneously slowed down the inactivation establishment (Eq. 8) and speeded up the inactivation removal (Eq. 7) of Nav1.7, as both rates are expected to be affected with the toxin (Maertens et al., 2006; Motin et al., 2016). Indeed, we checked that modifying only the establishment or the removal of the inactivation is enough to increase the spike frequency and amplitude derivative, as both effects add up. In the simulations of Fig. 3 and 4 we combined a 10 % variation in the values of both A parameters (Eq. 7 and 8).

**Fig 4.**
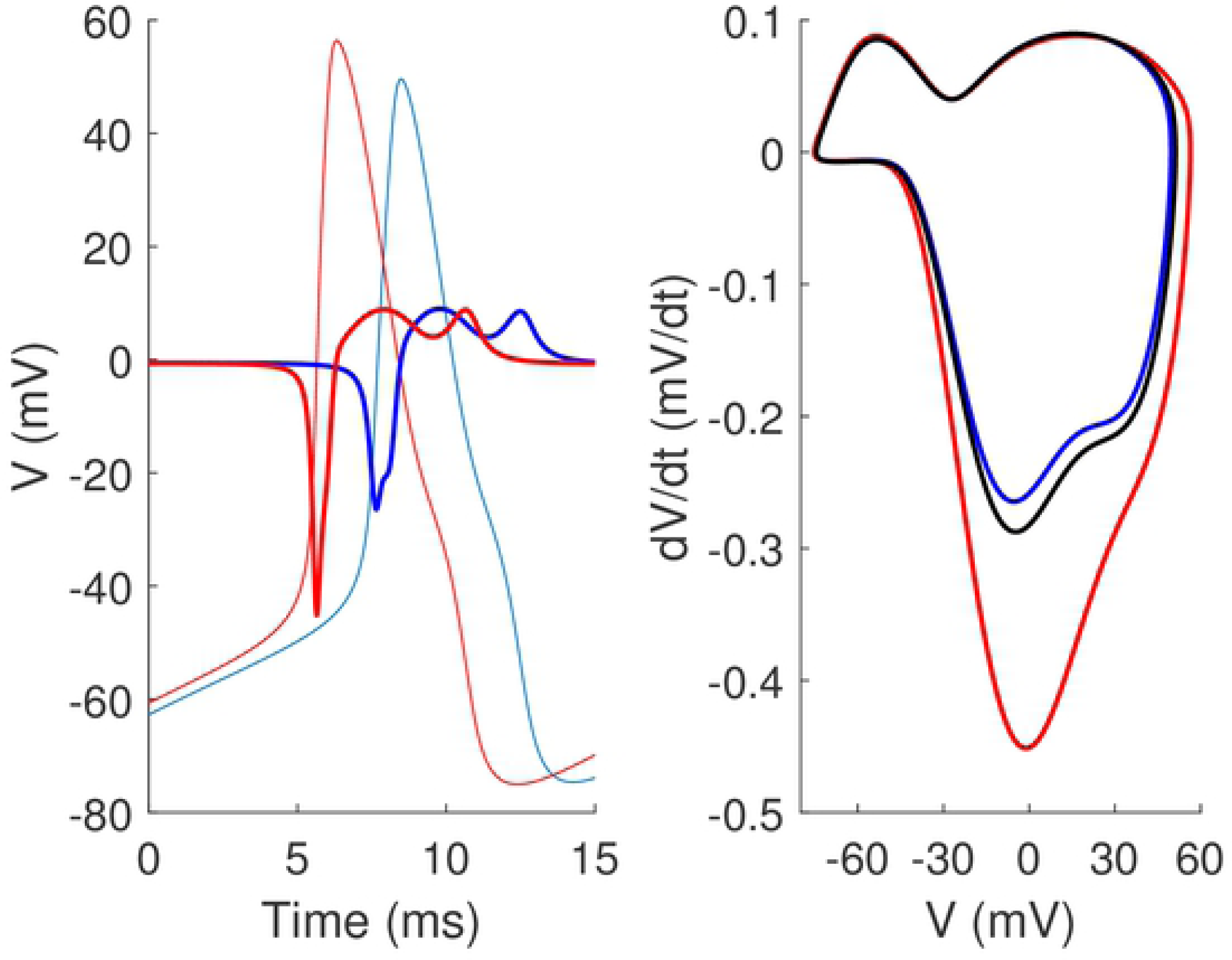
Simulations of membrane potential and its derivative. Third spike within a train of sustained spiking. In the left panel, the membrane potentials (mV) are plotted against time (ms) with thin lines and their corresponding minus derivatives (mV/dt) are plotted with thick lines (x100). Blue traces represent control conditions (A in a_h_ = 0.04 ms^−1^, A in b_h_ = 1.0 ms^−1^) and red traces represent OD1 application (A in a_h_ = 0.044 ms^−1^, A in b_h_ = 0.9 ms^−1^), both under a depolarizing current of 200 pA. In the right panel, the membrane potential (mV) is plotted against its minus first derivative (mV/dt), red line for OD1 and blue line for control. The black line shows the result for control gating parameters with a depolarizing current of 221.41 pA to match the spiking rate observed in the OD1 simulation.

Simulations lasted 120 ms, with step depolarisations starting at 10 ms and ending at 90 ms. Levels of constant current (pA) applied are specified in the corresponding figures. Note that the current levels needed to evoke single and multiple spiking, as well as the duration and amplitude of the spikes, are of the order to the values shown by Cao et al (2016) in current clamp recordings of hiPSC derived sensory neurons. Moreover, a recent paper using the same type of neurons reported a rheobase of 100 pA in healthy (control) conditions (see figure 6–A in McDermott et al., 2019), which is very close to the value produced by our model (138 pA). This confirms that the parameters are biologically plausible despite having been collected from publications using different methods and mammal species.

The model does not incorporate explicit temperature dependence through scaling factors (Q10), but it is based on equilibrium potentials and membrane properties (Choi and Waxman, 2011) as well as half activation voltages (Cummings and Waxman, 1997; Rush et al., 2007) measured at room temperature. Figures were created with Matlab (R2019a, The MathWorks, Inc.).

## Results

### Experimental recordings

We compared the spontaneous activity of sensory neurons obtained from a control subject (cell line AD3, n = 18 cultures) with an erythromelalgia patient (cell line RCi002-A, n = 24 cultures). The percentage of electrodes showing activity was 1.85 % in the control and 5.9 % in the disease cultures. After spike sorting we found 30 spontaneously active units in the patient and only 7 in the control. The firing frequency and amplitude of these units were not significantly larger in the disease cultures, implying that although much more units are active, they do not necessarily fire faster or have spikes of larger amplitude. It is worth mentioning that the fastest neurons and the tallest spikes were found in the disease cultures, but the distributions were not significantly different because slow units with small amplitude were present in both groups as well. The larger number of spontaneously active neurons found in the disease cultures is consistent with the results of a previous study (Cao et al., 2016).

The cultures from the control subject were treated with OD1 (100 nM), resulting in the percentage of active channels increasing from 1.85 to 6.02 % and a concomitant increase in the number of active units, from 7 to 24. An example of the voltage data before and after treatment is presented in Fig 1–A, note the increase in spontaneous activity rate and amplitude. The same effect was observed in an erythromelalgia culture conducted as a verification (not shown).

Pairing all the units that were active before and after OD1 application in the control cells (n = 7), we found that both the spike frequency and amplitude increased significantly (Fig 1–B). Although the increase in amplitude was not dramatic (20 % in average), it was consistently found in every pair of units. The extent of the increase in firing frequency was more pronounced (70 %). The unpaired units that were initially silent and started to fire after the OD1 were not significantly different in rate and amplitude from those already active before the treatment.

The cells from the patient were treated with PF05089771 (100 nM). The percentage of channels with spiking activity fell from 5.9 to 2.08 %. Twenty four out of thirty of the units sorted stopped firing in the presence of the drug. A typical example of the raw voltage trace before and after treatment is presented in Fig 2–A, note the suppression of the spiking activity. Pairing the few units that remained active after the treatment (n=6), we found that the spike frequency significantly decreased but the amplitude did not change (Fig 2–B). The unpaired units that stopped firing as a result of the PF05089771 application were not significantly different in rate and amplitude to the ones that remained active after treatment.

In order to assess for functional evidence of possible synaptic connectivity, we calculated cross correlation histograms between the firing times of the neurons that were recorded from the same electrode. No histogram peaks at fixed latency were found, which is compatible with the established notion that primary nociceptive neurons do not form synaptic contacts between them.

### Numerical simulations

In this subsection we present the results obtained with the computational model explained in Methods. The parameter set is provided in Table 1 and the model equations in S1 Appendix.

Erythromelalgia condition was simulated through a change in the Nav1.7 half-activation voltage parameter (*d* in Eq. 4) from −30 to −36 mV. OD1 effects were simulated through an increase in the constant that multiplies the inactivation removal rate (*A* from 0.04 to 0.044 in Eq. 7) and a simultaneous decrease in the constant that multiplies inactivation establishment rate (*A* from 1 to 0.9 in Eq. 8).

Fig. 3 shows simulations of control conditions (upper row), erythromelalgia (middle row) and OD1 treatment (bottom row). The minimum current level able to evoke a single spike (rheobase) is shown in the left column. The cyan traces are responses to constant currents just below threshold, while the blue traces show the rheobase responses. The current levels (printed in the corresponding panels) are of the order shown in Cao et al. (2018). Note that the rheobase decreased in both erythromelalgia and OD1 simulations, in comparison with the control.

The blue traces in the middle column of Fig. 3 show the minimum level of current needed to evoke sustained spiking. The current levels are printed in the corresponding panels, note again the decrease in erythromelalgia and OD1 simulations. The cyan traces represent responses to the highest level of current that evoked only a single spike.

The red traces of the third column show the response of control (upper row), erythromelalgia (middle row) and OD1 (bottom row) simulations to the minimum current level that was able to evoke continuous spiking in control conditions (146 pA). Note that both erythromelalgia and OD1 simulations reached higher spiking rate than control. The green plots show the same responses after blocking Nav1.7 receptors. The spiking activity stopped in all cases, mimicking the analgesic treatment with the Nav1.7 blocker PF-05089771 at a saturating dose.

The possible mechanism behind the enhanced extracellular spike amplitude observed after OD1 treatment (Fig. 1) is addressed in the simulation of Fig. 4. We focused on the third spike of the simulated train because it is representative of the continuous firing regime. The order of the spike is not important providing that we avoid the first one after the onset of the current step, which is slightly taller. This is due to less Na^+^ inactivation, as the first spike is the only one that starts from the resting potential before the stimulus. In the spikes that follow this initial transient, the peak value of the membrane voltage and Na^+^ currents reaches a stable level. By choosing the third spike of the train we make sure that our assessment applies to a typical spike within a train of repetitive firing, without being influenced by the initial conditions of the stimulus.

The simulation of OD1 treatment (A in a_h_ = 0.044 ms^−1^, A in b_h_ = 0.9 ms^−1^) is plotted with red traces and the control (A in a_h_ = 0.04 ms^−1^, A in b_h_ = 1.0 ms^−1^) with blue traces (Fig. 4), in both cases the model neuron spiked regularly in response to a 200 pA supra-threshold constant depolarizing current. Membrane voltages are represented with thin lines (Fig. 4, left panel). Note that the red spike (OD1) occurs before the blue spike (control), due to its higher firing rate (or shorter inter spike interval) with little amplitude difference. This picture changes when we look at the minus first derivative of the intracellular spike, which has been shown to constitute a reasonable estimation of the extracellular spike (e.g., Figure 1 in Henze et al., 2000). The thick traces in Fig. 4 (left panel) show the minus first derivative of the voltage (x100 for better graphical comparison with the voltage traces of the same colours). Note that the red derivative reached considerably larger amplitude than the blue one (thick traces), contrasting with the membrane voltages (thin traces) that showed small amplitude differences. The first derivative corresponds to the slope, meaning that the red spike reaches higher slope at its maximum point than the blue spike as a consequence of the Nav1.7 inactivation impairment.

In order to assess if any increase in the model firing rate is expected to come accompanied by an increase of the peak derivative, we created exactly the same frequency shift by increasing the level of constant current. The result is displayed in the simulation of the right panel of Figure 4, where we plotted the membrane voltage in abscissas against its minus first derivative in ordinates, for the third spike of the simulated train. Following the same colour code of the left panel, the simulation of OD1 is represented with red trace and the control with blue trace. The black line shows a simulation with control Nav1.7 inactivation rates (A in a_h_ = 0.04 ms^−1^, A in b_h_ = 1.0 ms^−1^) after increasing the constant current from 200 to 221.45 pA, in order to obtain exactly the same spike timing observed in the OD1 simulation. In this representation, the rightmost point of the orbits corresponds to the spike peak while their lowest point corresponds to the peak derivative. Note that the peak derivative reached larger value in the red than in the black plot, both exceeding the level observed in the blue plot. This means that increasing the spike rate through an impairment of Nav1.7 inactivation has a much stronger effect on the spike ascending slope (derivative) than the same rate shift created with an increased level of constant current. In contrast, the spike peak (right edge of the orbits) is the same in the red and black plots.

To summarize, we considered the hypothesis that the extracellular spike was taller after OD1 treatment mainly because the intracellular spike reached higher slope in the ascending phase. The model simulation of Fig. 4 confirmed the plausibility of the proposed mechanism and showed that the amplitude effect is expected to be larger when the rate increase is created through the inactivation parameters. However, our hypothesis cannot be truly confirmed without a simultaneous intracellular recording, which we aim to perform in the future.

## Discussion

In this study, we performed to our knowledge the first test of the scorpion toxin OD1 in hiPSC-derived sensory neurons recorded in MEAs. The main effect of this toxin is to block the fast inactivation process of the threshold current Nav1.7 (Motin et al., 2016). Our extracellular recordings also confirmed previous findings of Cao et al. (2016) with current clamp regarding the increased spontaneous firing of erythromelalgia sensory neurons and the “analgesic” *in vitro* effects of PF-05089771. The effects of both compounds were explained using a conductance-based computational model.

### Erythromelalgia and PF-05089771

Cao et al. (2016) performed intracellular recordings of hiPSC derived sensory neurons in the current clamp configuration, comparing erythromelalgia patients with control subjects. Despite the heterogeneity of the samples, they found a significantly higher proportion of spontaneously firing cells in patients compared to those from control donors. Moreover, the patient’s cells showed a lower rheobase and reached higher firing rates in response to stimulation with current steps of increasing amplitudes. These findings point to the existence of elevated excitability in erythromelalgia cells, a fact that we confirmed in the present study by showing higher prevalence of spontaneous activity in MEA recordings (see Results).

Different neuronal gain of function mutations causing erythromelalgia have in common the fact that the Nav1.7 channel opens with smaller depolarisations, with leftwards shifts in the current-voltage relationship ranging from 5 to 10 mV in the case of the F1449V mutation (Dib-Hajj et al., 2005). According to these reports, we adopted a plausible value of −6 mV in our schematic model simulations of the disease. After this manipulation, the rheobase and the threshold for sustained spiking decreased dramatically, driving the model neuron towards a spontaneous firing regime of higher frequency.

Spontaneous firing in our cultures is due only to intrinsic cell excitability, as there are no reports about the formation of synapses between primary sensory neurons *in vitro*. Accordingly, we did not find evidence of functional connections in cross correlation histograms between neuronal firing times. Furthermore, in dissociated cell cultures the development of a synaptic network is usually accompanied by the emergence of population bursts (Maeda et al., 1995) which we did not observe here. Glia cells are not present in the cultures, so their reported contribution to abnormal neuronal activity in intact DRG preparations *in vitro* (Belzer and Hanani, 2019) can also be ruled out.

Regarding the effects of Nav1.7 channel blockers, Cao et al. (2016) reported a dose dependent reduction in spontaneous firing and an increase in the action potential rheobase. We were able to supress the firing with a high dose of PF-05089771 (Fig 2), confirming the fact that the excitability of the cells plummets after the treatment. The clinical efficacy of the drug, however, was inconclusive in the study of Cao et al. (2016), showing statistical significance vs placebo only at the 10% level at the 4 to 5 and 8 to 9 hour time points after dosing. In other clinical reports the compound has failed to produce significant pain relief (Emery, 2016; McDonnell et al., 2018), contrasting with our *in vitro* recordings and numerical simulations. It has been suggested that the lack of penetration across the peripheral nerve sheet by hydrophobic molecules of high molecular weight can be behind the failure of PF-05089771 to produce significant pain relief when administrated systemically (Yekkirala et al., 2017).

### Effects of OD1

The excitability increase in the presence of OD1 reported with voltage clamp recordings (Maertens et al., 2006) was confirmed in our extracellular recordings by the increase in the proportion of electrodes showing spontaneous firing, together with the faster rate observed in the neurons that were already active before the toxin application. Remarkably, the toxin also kindled many silent neurons into activity. In addition, we found a consistent increase of the extracellular spike amplitude after OD1 which has not been described before with MEA recordings. We further discuss this last point in the next subsection with the numerical simulations.

Intra-plantar OD1 injections have been validated as a mouse model of NaV1.7-mediated pain, allowing Nav1.7 inhibitors profiling through measurement of neuronal firing and pain behaviours (Deuis et al., 2016). The effects of OD1 and analogue toxins were evaluated with voltage clamp whole cell recordings in Chinese hamster ovary (CHO) cells expressing human Nav1.7 channels (Deuis et al., 2016; Motin et al., 2016). These studies found that the decay phase of the current slows down in a concentration-dependent manner in the presence of the toxins due to impairment of fast inactivation. This effect, typical among α-scorpion toxins, is created by a combination of slower inactivation establishment with a faster recovery from inactivation (reprisal). β-scorpion toxins, on the other hand, shift the current activation to more negative potentials, as happens in erythromelalgia mutations. In the case of OD1 this last effect has small magnitude (Motin et al., 2016), in accordance with earlier observations in *Xenopus laevis* oocytes with Nav1.7 channels expression (Maertens et al., 2006). A minor effect on the voltage dependence of channel inactivation has been also reported at higher OD1 doses (300 mM, Deuis et al., 2016) than we used here (100 mM). Following these descriptions, we focused only on the inactivation to model OD1 effects in our computational model, slowing down the rate of inactivation establishment and speeding up its removal.

### Numerical simulations

The computational model that we present here is able to provide satisfactory explanations at a qualitative level for the main experimental findings of our study. We did not attempt to perform a complete exploration of the parameter space but all values are biologically plausible. In the absence of stimulation the membrane potential tends to its fixed point which is the resting potential. By injecting an amount of current of the order used in recordings of hiPSC derived sensory neurons (Cao et al., 2016; McDermott et al., 2019) the system reaches a critical point (e.g., Ermentrout and Terman, 2010) that corresponds to the spike threshold. In this very fast transition the stability of the system switches and a periodic solution arises, configuring what mathematicians in dynamical system theory call a “canard” (the French word for “duck”). In a “canard” (Börgers, 2017), large amplitude oscillations appear suddenly with very small variations of a driving parameter, the depolarizing current in our case (*I_ext_* in Eq. 15). At first the periodic oscillation comprises only one cycle (rheobase level). If the depolarizing current is sufficiently large the model generates a periodic response of repetitive spiking. In Fig. 3 we used these two very relevant thresholds (single spike and sustained repetitive firing) to illustrate the pain created by the enhanced firing of nociceptors that occurs in erythromelalgia patients (McDonnell et al., 2016), and in healthy individuals after a scorpion sting (e.g., Garfunkel et al., 2007). When the level of current is above the threshold for repetitive firing, the increased excitability is manifested with higher firing rate, which can be abolished by blocking the Nav1.7 receptors in the simulation, with no need of additional Nav1.8 targeting.

Regarding the increased amplitude of the extracellular spike observed after OD1, we suggest that it can be mainly due to an increased slope in the ascending phase of the intracellular spike, combined with a minor growth of its amplitude. This interpretation is based on the observation that the minus first derivative of the intracellular voltage is a reasonable approximation of the extracellular spike (Henze et al., 2000). We showed that the first derivative (slope) is far more sensitive to inactivation blocking than to increases of the depolarizing current (Fig. 4). The comparison is based on the fact that both manipulations created exactly the same effect on the spiking rate. Thus, although the intracellular spike is not much larger after OD1, the extracellular spike can grow dramatically because the intracellular slope is highly sensitive to the inactivation gating. Although the proposed mechanism is plausible, simultaneous intra and extracellular recordings are needed to confirm the hypothesis. A comparable scenario in the opposite direction has been described after treatment with the selective Nav1.8 blocking (e.g., Fig. 5 in Payne, et. al., 2015), in the sense that both the intracellular voltage and its derivative were decreased by the drug. An important difference, however, is that in our case the effect of the slope appears to be dominant.

One aspect that the model does not explain is that the few remaining spikes observed after PF-05089771 treatment showed no amplitude decrease, although the firing rate tended to decrease (p = 0.03 in 6 neurons). We checked that mild blocking of Nav1.7 conductance decreases the slope and thus the amplitude of the spike derivative in our model. The neurons that persisted in firing (Fig. 2) proved to be relatively resistant to a saturation dose of PF-05089771 that terminated the activity in 80 % of the units (24 out of 30). One possibility is that this subpopulation (20 %) was resilient because the neurons had less Nav1.7 expression. In this case, our model based on Nav1.7 effects would not be applicable. For OD1, all neurons active in control conditions were sensitive to the compound, both in the discharging rate and amplitude, suggesting that they had significant Nav1.7 expression.

### Extracellular recordings and ionic currents

Although extracellular recordings are typically used to detect firing times, they also have the theoretical ability to report about what are considered to be intracellular features of the action potential (Gold et al., 2006). We want to highlight this possibility of using extracellular recordings to detect changes in ionic currents. If researchers first evaluate them together with voltage clamp data for validation purposes, they can then benefit from the opportunity to observe more neurons simultaneously. Our computational model allowed to relate the amplitude of an extracellular spike with Nav1.7 gating. Whether this could really be the footprint of a current in the extracellular spike needs to be assessed with current clamp and whole cell voltage clamp recordings of hiPSC derived sensory neurons, designing the right experiments to have a direct measure all relevant model parameters.

### Model limitations

In this article we provide a starting point model to explain firing behaviour linked to chronic pain. Several of the relevant parameters were obtained from published observations in rats and humans (Table 1). In the absence of direct measurements, some parameters were filled with “educated guesses”, but they are biologically plausible. In order to keep the model simple and with a clear biological meaning and physical unit for all variables, we opted for a modified HH scheme, which also has the virtue of being familiar to neurobiologists although more refined kinetic schemes have been introduced (e.g., Andreozzi et al., 2019). We included only one potassium current of non-inactivating type to avoid unnecessary complexity. The only added current to the original HH scheme was the Nav1.7, because it is strictly needed to implement the biological mechanisms under consideration. A detailed quantitative modelling would require Ca^++^ currents (e.g., Scroggs and Fox, 1992) and several types of Na^+^ (Rush et al., 2007; Zhang et al., 2017) and K^+^ (Zemel et al., 2018) currents that have been found in mammalian DRG cells, and is beyond the scope of this study. We are adopting a level of simplification in the modelling process that is fitted for the purpose of explaining our experimental results with MEA recordings regarding compounds that specifically target Nav1.7 receptors.

### Importance of the interaction between sodium currents

The main focus of this discussion is Nav1.7 due to its direct relation with the compounds tested. OD1 is a potent and selective modulator of NaV1.7, which does not affect Nav1.8 (Maertens et al., 2006) and PF-05089771 is a selective blocker of Nav1.7 receptors (Cao et al., 2016). However, we must take into account the complex interaction between Nav1.7 and Nav1.8 in the excitability tuning of sensory neurons (Choi and Waxman, 2011). Indeed, modulations of Nav1.8 alone largely affect the excitability. For example, the selective Nav1.8 channel blocker PF-01247324 attenuates neuronal firing and reduces the slope and peak value of the action potential in dissociated rat and human DRG neurons from organ donors with short post-mortem delay (Payne, et. al., 2015).

To resolve the conundrum of how Nav1.7 and Nav1.8 can be successfully targeted to treat neuropathic and inflammatory pain is beyond the scope of this article, but we want to finish this discussion summarizing a remarkable example (Rush et al., 2006, and references therein) of how the interaction between sodium currents can have profound functional consequences for the clinical signs of erythromelalgia. As explained in the Introduction, in pain pathways Nav1.7 increases neuronal excitability mainly by amplifying the depolarization caused by peripheral stimuli, allowing to trigger Nav1.8 which then creates most of the spike. This facilitates firing and pain, an effect that is exaggerated in erythromelalgia patients. A key property for this firing to occur (as implemented in our model) is that Nav1.8 does not inactivate until the voltage is quite high (e.g., −30 mV). In the sympathetic nervous system the situation is different because there are no Nav1.8 channels and spikes are sustained by other sodium currents that inactivate at lower voltage (Rush et al., 2006). Then, the excessive depolarization created by a mutated Nav1.7 in erythromelalgia can be sufficient to trigger inactivation, terminating the firing in the sympathetic nerves that mediate the contraction of peripheral blood vessels. This results in excessive vasodilatation, creating the typical sign of red extremities that gave name to the disease (erythromelalgia could be translated as “red neuralgia of the extremities”). The importance of the interaction between sodium currents is asserted by this example, because the same Nav1.7 neuronal mutation produces opposite effects in the firing of pain afferents and autonomic motor neurons.

### Concluding remarks

In this study we first used MEAs to evaluate Nav1.7 gating in hiPSC derived sensory neurons. Previous voltage clamp reports regarding the excitability effects of OD1 and PF-05089771 were confirmed. In addition, we found an unexpected effect of OD1 on the extracellular spike amplitude. To explain the results we constructed a conductance based computational model that highlights the importance of the intracellular spike slope to determine the amplitude of the extracellular spike.

## Acknowledgements

Study funded by MRC BH171892.

